# Investigate c-Fos changes in genetically identified amygdala neurons after mild footshock stress

**DOI:** 10.1101/2024.11.03.621553

**Authors:** Hallie Dong, Oliver M. Schlüter, Xiaojie Huang

**Affiliations:** Sewickley Academy, 315 Academy Avenue, Sewickley, PA 15143; Department of Neuroscience, University of Pittsburgh, Pittsburgh, PA 15260

**Author notes:** Corresponding Authors: Dr. Xiaojie Huang Corresponding Address: Dept. Neuroscience, Univ. of Pittsburgh, A210 Langley Hall / 5^th^ & Ruskin Ave, Pittsburgh, PA 15260. Competing financial interest: The authors declare no competing financial interest. Author contributions: H.R.D., O.M.S., and X.H. designed the experiments and analyses; H.R.D. and X.H. conducted the experiments. H.R.D. and X.H. conducted data analysis; H.R.D., O.M.S., and X.H. wrote the manuscript.

**Keywords:** morphine, cocaine, accumbens, silent synapse, D1 receptor, D2 receptor

## Abstract

The amygdala is a key brain region that processes stress-related inputs to reshape future behaviors. The lateral amygdala (LA), basolateral amygdala (BLA), and central amygdala (CeA) are important subregions that mediate different aspects of stress experiences from receiving sensory input to memory formation and behavioral responding. The principal neurons in these regions are glutamatergic pyramidal neurons, which are genetically separable into two subpopulations, protein phosphatase 1 regulatory subunit 1B-positive (Ppp1r1b, also known as DARPP-32) parvocellular neurons and R-spondin2-positive (Rspo2) magnocellular neurons. Recent studies show that these two subpopulations of amygdala neurons differentially regulate appetitive versus aversive behaviors. The research goal of this study is to explore whether amygdala Ppp1r1b and Rspo2 neurons are transcriptionally activated by moderate stress experience, such that persistent cellular changes are made to influence future functional output of these two subtypes of neurons. To test transcriptional activation, we focused on c-Fos, one of the early genes that are transiently expressed in response to cellular stimulations to regulate downstream gene transcription. Moderate stress was introduced through brief footshocks, with mice without footshock as controls. Between shocked and control mice, we observed similar numbers of Ppp1r1b or Rspo2 neurons per unit area that expressed c-Fos, which was consistent across LA-BLA and CeA. Moreover, in LA-BLA, Ppp1r1b/c-Fos cells consistently outnumber Rspo2/c-Fos cells across treatment conditions, and the reverse is true in CeA. These results suggest that moderate stress experience is not sufficient to induce robust transcriptional alterations in the two key subpopulations of amygdala neurons, and Ppp1r1b versus Rspo2 neuron activities, as measured by c-Fos expression levels, show differential dominance in amygdala subregions.

## Introduction

Stress is a common trigger for many psychiatric disorders (Davis et al., 2017; Schneiderman et al., 2005). Strong and persistent stress experience induces profound molecular and cellular alterations in various brain regions that regulate emotion and motivation (McEwen et al., 2016). A critical such brain region is the amygdala, where glutamatergic principal neurons respond to a variety of stress-related experience, such as drugs of abuse (Kufahl et al., 2009), social stress (Nosjean & Granon, 2022), restraint stress, injury/pain-related stress, and fear (Rajbhandari et al., 2016). The amygdala subregions, including the lateral amygdala (LA), basal lateral amygdala (BLA), and central amygdala (CeA), contribute to different aspects of processing the emotional experience. For example, LA receives and transmits external sensory inputs (Yang & Wang, 2017), BLA develops conditional fear and fear memory (Gale et al., 2004), and CeA produces behavioral and physiological responses (Gilpin et al., 2015).

Stress-induced cellular changes can be detected by the expression of c-Fos (Hudson, 2018), an immediate early gene rapidly expressed in response to cellular stimulations, which subsequently regulates downstream gene transcription. Electric footshock is an acute stressor widely used in rodent research to induce pathophysiological emotional and motivational conditions (Pacak & McCarty, 2007). However, results about the impact of footshock on c-Fos expression in the amygdala are not consistent. Specifically, while some studies show an overall increase in mRNA levels of c-*fos* in the amygdala as a whole (Campeau et al., 1991), other studies do not detect such a change (Lin et al., 2018; Smith et al., 1992). In studies in which footshock-induced c-Fos increases are observed in the amygdala, the effects are often confined within subdivisions of the lateral, medial, or central amygdala (Lanuza et al., 2008; Milanovic et al., 1998; Pezzone et al., 1992). These inconsistent results raise a possibility that different subregions and/or subtypes of amygdala neurons respond to footshock stress differently.

Principal neurons in the amygdala are genetically separable into two subpopulations, protein phosphatase 1 regulatory subunit 1B-positive (Ppp1r1b, also known as DARPP-32) parvocellular neurons (Kim et al., 2017) and R-spondin2-positive (Rspo2) magnocellular neurons. These two subpopulations of neurons respond to appetitive versus aversive stimuli by differentially expressing c-Fos, and regulate reward seeking versus aversion avoidance distinctively (Kim et al., 2017). To examine whether these two amygdala neuronal subpopulations respond to footshock stress differently, we employed two transgenic mouse lines, in which Ppp1r1b or Rspo2 neurons are tagged with yellow fluorescence, such that they can be detected visually. We treated the mice with a moderate footshock procedure, and quantified c-Fos expression in Ppp1r1b or Rspo2 neurons across amygdala subregions. We detected basal expressions of c-Fos in both Ppp1r1b and Rspo2-positive neurons in LA-BLA and CeA, presumably in response to new environment. Following footshock experience, the density of c-Fos-expressing Ppp1r1b neurons or c-Fos-expressing Rspo2 neurons did not change in LA-BLA or CeA. Moreover, Ppp1r1b/c-Fos cell densities were consistently higher than Rspo2/c-Fos cells in LA-BLA, whereas Ppp1r1b/c-Fos cell densities were lower than Rspo2/c-Fos cells in the CeA. These results suggest that 1) the two major subpopulations of amygdala principal neurons are not highly responsive to acute footshock; and 2) depending on the specific amygdala subregions, Ppp1r1b and Rspo2 neurons exhibit differential dominance in their population activities.

## Results

### Generation of mice in which Ppp1r1b and Rspo2 neurons are fluorescently tagged

Both Cartpt-Cre and Rspo2-Cre mice with the C57BL/6J background were obtained from the Tonegawa lab (Kim et al., 2016). We crossed these mice with R26-*loxP*-STOP-*loxP*-EYFP mice, such that enhanced yellow fluorescent protein (EYFP) expression was selectively detected in neurons expressing Cre under Cartpt- or Rspo2-promoters (Fig. 1A). The Ppp1r1b-Cre mouse line is not available. However, Ppp1r1b and Cartpt are co-expressed exclusively (Kim et al., 2016). As such, Ppp1r1b neurons can be experimentally accessed by targeting Cartpt.

**Figure 1.**
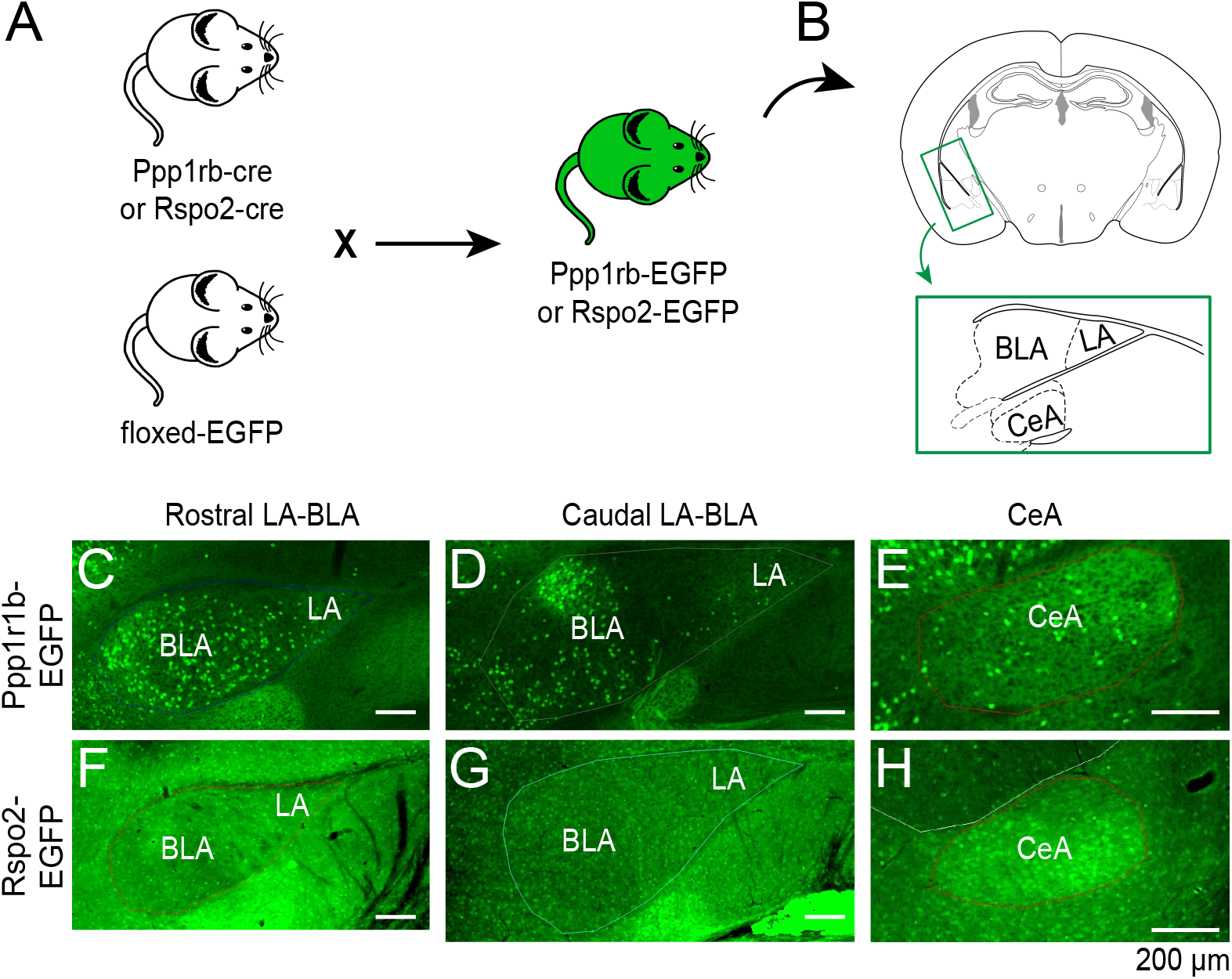
Generation of Ppp1r1b-EYFP and Rspo2-EYFP transgenic mouse lines. **A** Diagram for generating the two mouse lines. **B** Diagram of a coronal section containing amygdala subregions. **C**-**E** Ppp1r1b-EYFP expression in rostral LA-BLA (**C**), caudal LA-BLA (**D**), and CeA (**E**). Ppp1r1b-EYFP cells were evenly distributed in the rostral LA-BLA (**C**), appeared in patches in the caudal LA-BLA (**D**), and in low densities in CeA (**E**). **F-H** Rspo2-EYFP expression in rostral LA-BLA (**F**), caudal LA-BLA (**G**), and CeA (**H**). Rspo2-EYFP cells were evenly distributed in low densities in the rostral (**F**) and caudal LA-BLA (**G**), and in high densities in CeA (**H**). Scale bars = 200 µm

### Ppp1r1b and Rspo2 neurons in amygdala subregions

We prepared the brain slices that contained the amygdala at a thickness of 50 µm. After fixation and staining for EYFP, we imaged LA-BLA and CeA neurons under the fluorescence microscope. In Ppp1r1b-Cre mice, EYFP-expressing neurons were evenly distributed in rostral LA-BLA (Fig. 1A) and appeared in patches in caudal LA-BLA (Fig. 1B). There was also a low density of EYFP-expressing cells in the CeA (Fig. 1C). In Rspo2-Cre mice, EYFP expression appeared evenly at low levels in the rostral (Fig. 1D) and caudal LA-BLA (Fig. 1E), and high levels in the CeA (Fig. 1F).

### c-Fos in LA-BLA Ppp1r1b and Rspo2 neurons after footshock

To test how Ppp1r1b and Rspo2 neurons may respond to footshock, we harvested the brains 90 min following the footshock test from mice undergoing footshocks versus mice exposed to a similar chamber but without the footshocks. We then stained the tissue containing the amygdala for the immediate early gene c-Fos as an indicator of recent intense neuronal activity (Hudson, 2018). In LA-BLA, the overall c-Fos expression did not change following footshock (*t*(16)=0.393, *p* = 0.700, unpaired *t*-test, Fig. 2A-C). Moreover, neither Ppp1r1b nor Rspo2 cells exhibited differential c-Fos expression following footshock (genotype x treatment interaction: F(1, 13) = 1.929, *p* = 0.188, two-way ANOVA; Fig. 2A, B, D). However, Ppp1r1b cells showed an overall higher expression of c-Fos than Rspo2 cells (main effect of genotype: F (1, 13) = 8.256, *p* = 0.013, two-way ANOVA, Fig. 2D; *t*(15)=3.177, *p* = 0.006, unpaired *t*-test, Fig. 2E). These results suggest that in LA-BLA, there are more Ppp1r1b than Rspo2 cells that express c-Fos under basal conditions or upon exposure to a novel environment, although neither cell types show further elevated c-Fos expression to footshock stimulations.

**Figure 2.**
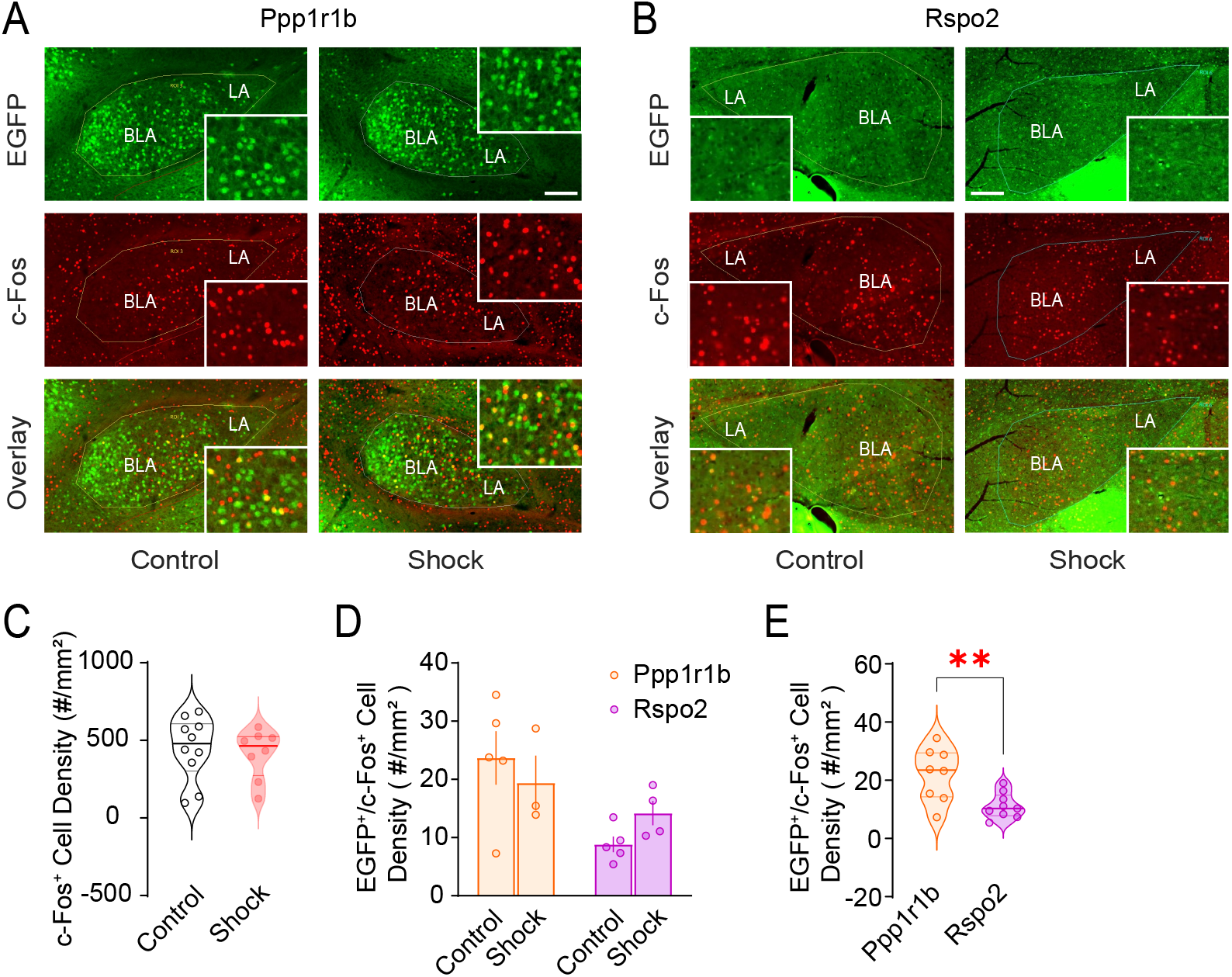
c-Fos expression in LA-BLA Ppp1r1b and Rspo2 cells without and with footshock. **A** A subpopulation of LA-BLA Ppp1r1b cells co-express c-Fos without or with footshock experience. **B** A subpopulation of LA-BLA Rspo2 cells co-express c-Fos without or with footshock experience. **C** Overall c-Fos+ cell densities in LA-BLA did not change following footshock. **D** c-Fos expression in LA-BLA Ppp1r1b or Rspo2 cells did not change following footshock. **E** LA-BLA Ppp1r1b cells co-express c-Fos more than Rspo2 cells across treatment conditions (shock or no-shock combined). Scale bars = 200 µm for low-magnifications and 100 µm for high-magnification inserts. ** *p*<0.01

### c-Fos in CeA Ppp1r1b and Rspo2 cells after footshock

Beyond the LA-BLA, how may CeA Ppp1r1b or Rspo2 cells respond to footshocks? We first analyzed overall c-Fos expression in the CeA, which did not change significantly following the footshocks (*t*(15)=0.8151, *p* = 0.428, unpaired *t*-test, Fig. 3A-C). We found that CeA subpopulations of both Ppp1r1b and Rspo2 cells co-expressed c-Fos in the absence or presence of footshocks (genotype x treatment interaction: F(1, 13) = 2.531, *p* = 0.136, two-way ANOVA; Fig. 3A, B, D), whereas Ppp1r1b cells showed an overall lower % expression of c-Fos than Rspo2 cells (main effect of genotype: F (1, 13) = 13.35, *p* = 0.003, two-way ANOVA, Fig. 3D; *t*(15)=3.595, *p* = 0.003, unpaired *t*-test, Fig. 3E). Finally, both c-Fos+ Ppp1r1b cells and c-Fos+ Rspo2 cells were small fractions of the total c-Fos+ cells (comparing Fig. 3E to Fig. 3C). Together, these results suggest that in the CeA, there are less Ppp1r1b cells than Rspo2 cells that express c-Fos upon exposure to a novel environment, although neither cell types show further elevated c-Fos expression to footshock stimulations.

**Figure 3.**
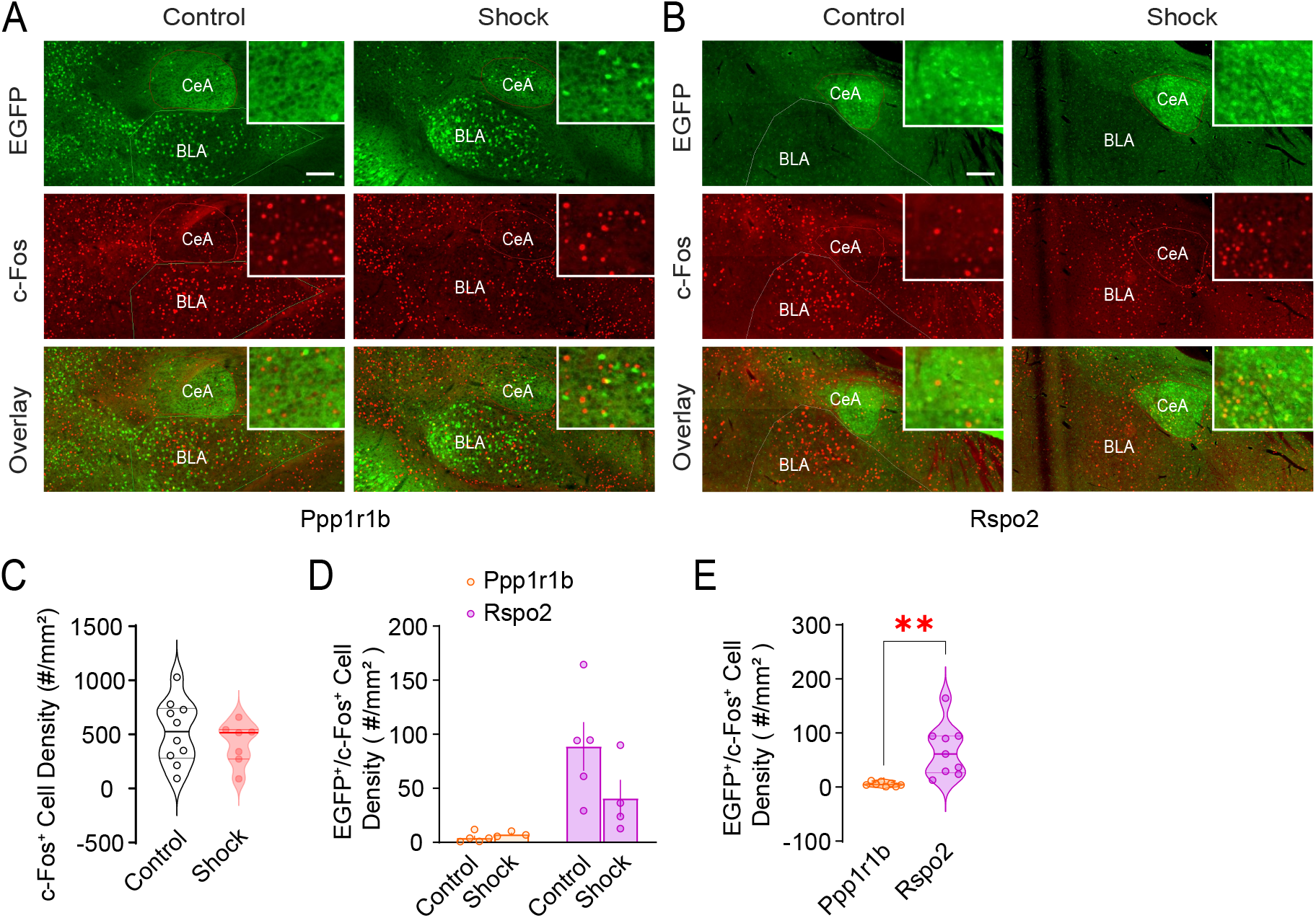
c-Fos expression in CeA Ppp1r1b and Rspo2 cells without and with footshock. **A** A subpopulation of CeA Ppp1r1b cells co-express c-Fos without or with footshock experience. **B** A subpopulation of CeA Rspo2 cells co-express c-Fos without or with footshock experience. **C** Overall c-Fos+ cell densities in CeA did not change following footshock. **D** c-Fos expression in CeA Ppp1r1b or Rspo2 cells did not change following footshock. **E** CeA Ppp1r1b cells co-express c-Fos less than Rspo2 cells across treatment conditions (shock or no-shock combined). Scale bars = 200 µm for low-magnifications and 67 µm for high-magnification inserts. ** *p*<0.01

## Discussion

Following footshock versus no shock we did not detect changes in c-Fos+ cell densities in LA-BLA or CeA. While the CeA results are consistent with previous report – that CeA c-Fos expression does not change following exposure to aversive stimuli such as foot shocks, novel environment, restraint, or air puffs (Knapska et al., 2007), the LA-BLA c-Fos results were not expected. In particular, *Rspo2* was shown to be enriched in footshock-responding BLA neurons (Kim, Tonegawa, 2016), thus it was expected that following footshock there should be an increase in Rspo2/c-Fos cells in BLA. It is likely that even in no-shock mice, exposure to a novel environment – the testing chamber, and handling of the mice effectively increase expression of immediate early genes in LA-BLA (Gouty-Colomer et al., 2016), rendering low signal-to-noise ratio for detecting footshock-induced c-Fos increase on top of the elevated basal level. Current study allowed the mice to adapt to the testing chamber for 20 min on the testing day before footshocks, which may not be sufficient to lower the background expression of c-Fos. Indeed, a previous study in rats allowed 10 days for the rats to adapt to handling and the testing chamber before footshocks were applied on the 11th day (Campeau et al., 1991). A similar study in mice allowed for 3 days of experimenter handling before footshocks (Kim et al., 2016). These repeated handling before footshocks presumably minimize background c-Fos expressions. Moreover, the electric shocks we used (3 × 2-sec pulses at 0.75 mA repeated at 2 min intervals) were mild, as compared to shocks at 0.6 mA for 0.5 sec x 10 pulses per train x 5 trains – which were shown to increase amygdala c-*fos* mRNA expression in rats (Campeau et al., 1991). Nonetheless, shocks at 0.75 mA for 2 sec x 3 pulses were reported to increase amygdala immediate early gene Arc expression in mice (Gouty-Colomer et al., 2016). Finally, our detection sensitivity may be further enhanced by sampling more specific subregions of the amygdala, e.g. differentiate along the rostral-caudal axis (Kim et al., 2016), and possibly using different immediate early gene markers such as Arc (Bonapersona et al., 2022; Gouty-Colomer et al., 2016).

Under current procedures, we did not detect changes in c-Fos expressions in Ppp1r1b or Rspo2 subpopulations of neurons following footshocks. Nonetheless, both populations contribute to c-Fos expression without or with footshocks (Figs. 2A,B,D; 3A,B,D). Moreover, the two populations exhibit different c-Fos expressions in amygdala subregions – in LA-BLA, Ppp1r1b neurons show more c-Fos expression than Rspo2 neurons without or with footshocks, and the reverse is true in CeA. Thus, either when the mouse was adapting to the novel environment or experiencing footshocks, LA-BLA information processing recruits more Ppp1r1b than Rspo2 neurons, whereas the CeA engages more of Rspo2 neuron activities.

## Materials and methods

### Animals

Cartpt-EYFP and Rspo2-EYFP mice were generated by breeding Cartpt-Cre or Rspo2-Cre mice with R26-*loxP*-STOP-*loxP*-EYFP mice (JAX 006148) respectively (Fig. 1A). Both male and female mice were used at the age of postnatal day 30-43. Mice were group-housed at 22□±□1□°C and controlled humidity (60□±□5%), on regular 12:12 light-dark cycles. Footshocks were performed during the light phase. All animal use was in accordance with protocols approved by the Institutional Animal Care and Use Committees at the University of Pittsburgh.

### Footshocks

Both male and female mice of either genotypes were randomly assigned to the control or shock groups. The mice were placed in a fear conditioning chamber (Med Associates) and allowed to freely explore the chamber for 20 minutes. At the end of 20 min, mice in the shock group were given 3 shocks, 120 seconds apart, at an intensity of 0.75 mA for 2 sec. Mice in the control group did not get shocks.

### c-Fos and EYFP immunohistochemistry

90-120 minutes after the footshocks, the mice were overdosed with isoflurane and transcardially perfused with ice-cold 0.1 M phosphate buffer (PB) followed by ice-cold 4% paraformaldehyde in PB. The brains were then removed and placed in 4% paraformaldehyde at 4°C for 24-48 hours. The brain tissue containing the amygdala was then sliced into coronal sections of 50 µm on a vibratome (Leica VT 1200s). The sections were immersed in 0.5% sodium borohydride in 0.1 M PB, then 0.15% hydrogen peroxide in 0.1 M PB, then incubated at 4°C for 1-3 days in the presence of Triton X-100 (0.3%, Sigma), 1% normal donkey serum (Jackson ImmunoResearch, West Grove, PA), rabbit-anti-c-Fos antibody (1:1000; Cell Signaling Technology #5348, Danvers, MA), and goat-anti-YFPYFP antibody (1:500; MyBioSource #MBS448123, San Diego, CA). The next day, tissues were incubated in secondary antiserum with donkey-anti-rabbit (Invitrogen, Alexa Fluor 594, A-21207, 1:500) and donkey-anti-goat (Invitrogen, Alexa Fluor 488, A-11055, 1:500) for 2 hours in the presence of Triton X-100 (0.3%) and 1% normal donkey serum. The sections were then mounted onto superfrost microscope slides (Fisherbrand) using Fluoromount-G mounting medium (Life Technologies Corp., Carlsbad, CA), left at room temperature overnight, then sealed and stored in 4°C.

### Imaging and c-Fos counting

Fluorescence images were taken at 10x magnification using a scanning microscope (Olympus Slideview VS200), and analyzed using Olympus CellSens software. We quantified ∼15-20 regions of interest (ROI) per animal for LA-BLA subregions and ∼5 ROI per animal for CeA, and counted all sections manually. All counts were repeated 1-3 times to ensure accuracy and blinded to the treatment conditions. c-Fos+ cell data was converted to density (number of c-Fos+ cells / µm^2^), then averaged for each mouse for statistical analysis.

### Statistics

Individual mouse data was grouped by genotype or treatment and was analyzed using two-way ANOVA or unpaired *t*-test. All results are shown as mean±SEM.

## Acknowledgement

We thank Li Cai for technical support.

